# Time-of-day-dependent variation of the hepatic transcriptome and metabolome is disrupted in non-alcoholic fatty liver disease patients

**DOI:** 10.1101/2023.04.05.535334

**Authors:** Manuel Johanns, Joel T. Haas, Violetta Raverdy, Jimmy Vandel, Julie Chevalier-Dubois, Loic Guille, Bruno Derudas, Benjamin Legendre, Robert Caiazzo, Helene Verkindt, Viviane Gnemmi, Emmanuelle Leteurtre, Mehdi Derhourhi, Amélie Bonnefond, Philippe Froguel, Jérôme Eeckhoute, Guillaume Lassailly, Philippe Mathurin, François Pattou, Bart Staels, Philippe Lefebvre

## Abstract

Liver homeostasis is ensured in part by time-of-day-dependent processes, many of them being paced by the molecular circadian clock. Liver functions are compromised in non-alcoholic fatty liver (NAFL) and non-alcoholic steatohepatitis (NASH), and clock disruption increases susceptibility to non-alcoholic fatty liver disease (NAFLD) progression in rodent models. We therefore investigated whether time-of-day-dependent transcriptome and metabolome are significantly altered in human NAFL and NASH livers. Liver biopsies, collected within an 8 hour-window from a carefully phenotyped cohort of 290 patients and histologically diagnosed to be either normal, NAFL or NASH hepatic tissues, were analyzed by RNA sequencing and unbiased metabolomic approaches. Time-of-day-dependent gene expression patterns and metabolomes were identified and compared between histologically normal, NAFL and NASH livers. We provide here a first-of-its-kind report of a daytime-resolved human liver transcriptome-metabolome and associated alterations in NAFLD. Transcriptomic analysis showed a robustness of core molecular clock components in NAFL and NASH livers. It also revealed stage-specific, time-of-day-dependent alterations of hundreds of transcripts involved in cell-to-cell communication, intra- cellular signaling and metabolism. Similarly, rhythmic amino acid and lipid metabolomes were affected in pathological livers. Both TNFa and PPARγ signaling are predicted as important contributors to altered rhythmicity. NAFLD progression to NASH perturbs time-of-day-dependent processes in human livers, while core molecular clock component differential expression is maintained.

## INTRODUCTION

Normal tissue homeostasis requires precisely timed expression of proteins, including metabolic enzymes, around the clock and in accordance with entrained rhythms of light exposure, feeding periods and physical activity. The central clock, located in the suprachiasmatic nucleus, is light-entrained and connects to peripheral tissues to synchronize clock oscillators in these tissues. It is now clear that peripheral tissue clocks, including the hepatic clock, can operate autonomously *i.e.* independently of the light-entrained central hypothalamic clock and are not necessarily aligned. Indeed, the major *Zeitgeber* (“time giver”) setting the liver clock is food intake/nutrient availability rather than daylight, though the systems are mechanistically similar (*1, 2*). Much of what we know about circadian regulation derives from studying mice (*3*), which are nocturnal by nature. A global picture emerged from these studies, defining a molecular clock machinery. This cell-autonomous circadian core clock (CC) is made of 2 autoregulatory loops comprising 14 transcription factors, encoded by so-called core clock genes (CCGs). The first transcriptional loop is composed of the heterodimeric BMAL1 (*ARNTL*) and CLOCK (or alternatively NPAS2) transcriptional activators and of the PER and CRY transcriptional repressors. The secondary loop is constituted by the nuclear receptors RORs, REV-ERBα and REV-ERBβ. These interlocked transcriptional-translational feedback loops (TTFLs) define a cell-autonomous clock machinery which controls clock output genes (*4*), in turn regulating multiple cellular functions (*3*).

Because most primates (including humans) are diurnal, there are likely important differences from rodents in circadian regulation that have yet to be fully explored. A limited number of human time-of-day-resolved transcriptomes is available, especially for internal organs. Indeed, most studies undertaken with human biological material over a variable range of time-of-day points analyzed either whole blood (*5, 6*), peripheral blood mononuclear cells (PBMCs) (*7*), skin epidermis (***8***), subcutaneous white adipose tissue (*9, 10*), heart (*11*) or skeletal muscle (*12–14*) in a relatively low number of subjects (less than 30). A recent study investigated gene expression in subcutaneous white adipose tissue and skin from 625 healthy volunteers, allowing the identification of common, time-of-day-regulated genes which were strongly enriched in CCGs (*15*).

It is now clear from multiple experimental models in various species that circadian rhythm synchrony, which is critical for metabolic homeostasis, is deregulated at both molecular and functional levels in various disease states, especially in obesity and its complications like non-alcoholic fatty liver disease (NAFLD). NAFLD is a spectrum of liver conditions characterized by hepatic steatosis which combines, upon time, with varying degrees of necroinflammation. In its more severe, but still asymptomatic form, called non-alcoholic steatohepatitis (NASH), there is a significant risk if left untreated to evolve toward liver fibrosis and eventually cirrhosis and hepatocellular carcinoma, as well as cardiovascular diseases (*16*). Several large-scale NAFLD patient cohorts have now been analyzed focusing on differences in gene expression between disease stages (*17–20*). However, these datasets did not provide information about the timing of biopsies, while rhythmic processes and timed gene expression in the liver are reported to be deregulated in chemically-induced liver fibrosis or after high-fat feeding in rodents (*21, 22*). Whether and how deregulation of rhythmic biological processes occurs in human liver in the context of NAFLD progression remains totally unknown. Mure *et al* (*23*) recently established a 24 hour-circadian transcriptome atlas of 64 tissues from healthy baboons (*Papio anubis*). Relatively few robustly cycling genes were identified in the liver and CCGs were surprisingly not amongst these. Ruben *et al* (***24***) applied an algorithm (CYCLOPS) to infer rhythmic gene expression patterns of tissues collected *post mortem* from human donors. Though this study included over 600 donors, only 648 genes were predicted to exhibit time-of-day-dependent expression (FDR<0.05) using this method, and only a few known CCGs were identified. Thus, both ethical and technical hurdles are a major impediment to fully appreciate time-of-day dependency of gene expression in healthy liver and its deregulation in NAFLD. Importantly, a similar conclusion can be reached when considering the hepatic metabolome, which is clock-controlled in mice (*22, 25, 26*) and disturbed in various liver dysfunction models (*22, 27, 28*). Disturbances in the hepatic chronometabolome observed in rodent NAFLD models have not been reported for humans so far (*29*).

Considering these knowledge gaps, we therefore asked whether the hepatic time-of-day-dependent transcriptome and metabolome are affected during NAFLD progression. We leveraged a large cohort of morbidly obese patients undergoing bariatric surgery from whom liver biopsies were taken peri-operatively [HUL cohort (*18*)]. Hepatic transcriptomes and metabolomes were obtained from a sub-cohort of 318 patients histologically diagnosed as definite NAFL, NASH or normal livers. Amongst these, the exact biopsy time-of-day was known for 290 patients (Figure 1A). Using this information in an original approach integrating multiple statistical tests, we provide the first-ever robust analysis of time-of-day-dependent gene expression and tissue metabolite abundance in human liver, as well as changes associated with the different stages of NAFLD.

**Figure 1:**
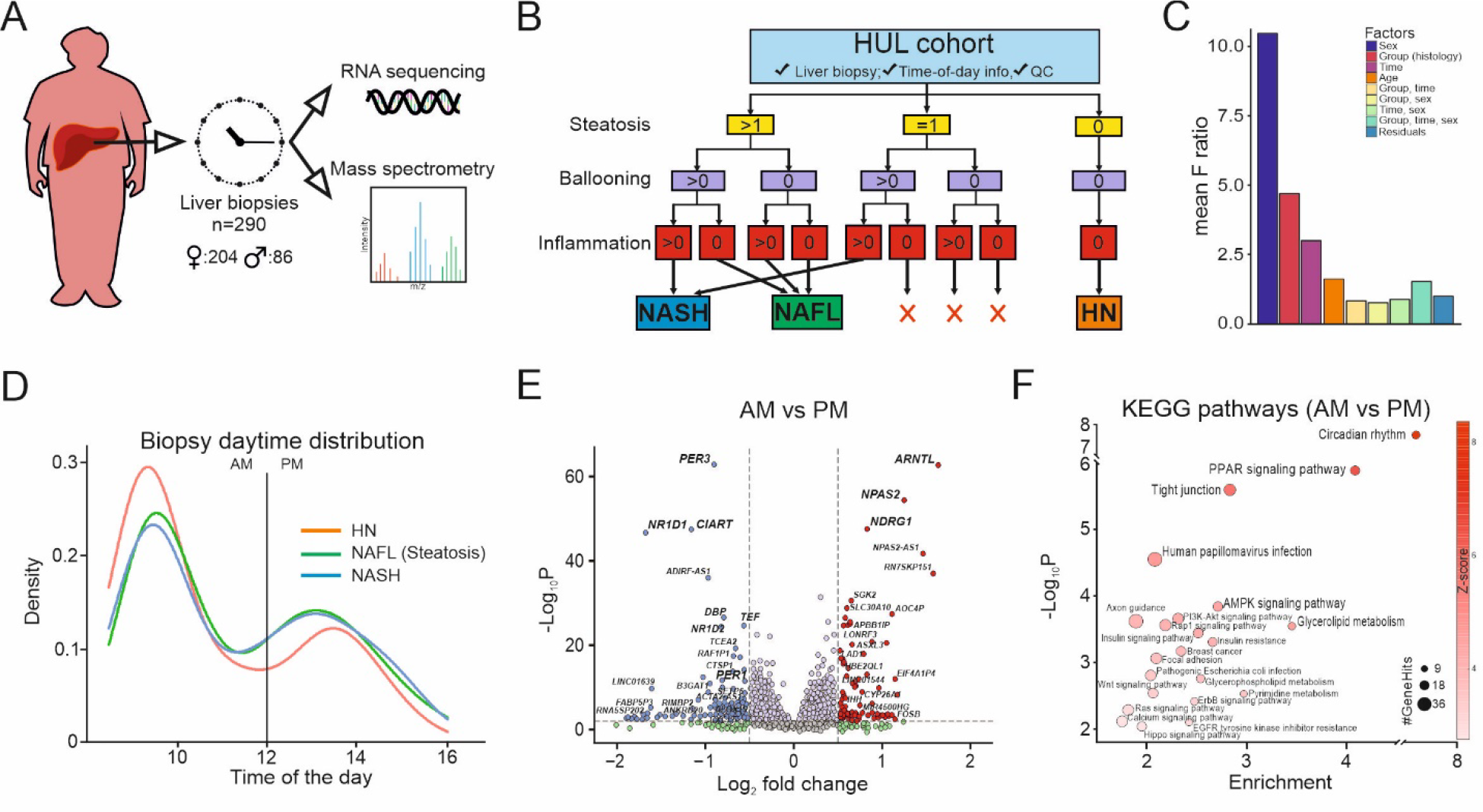
**Timed liver biopsies from a large cohort of humans with obesity. A:** Liver biopsies with available exact daytime information were obtained from 290 morbidly obese (BMI>35) patients (204 women, 86 men) while undergoing bariatric surgery. Transcriptomes were obtained by bulk RNA sequencing and metabolomes by multimode LC-MS. B: To determine whether the disease state of the liver alters circadian gene expression, biopsies were classified into three groups: “HO” (healthy obese), “NAFL” (benign steatosis only) or “NASH” (steatosis+inflammation) following the indicated decision tree. **C:** ANOVA was used to reveal the main sources of overall inter-sample variation in gene expression, highlighting sex, liver disease state and daytime of biopsy collection as main sources of variation. **D:** Liver biopsy collection daytimes range from 8 AM to 4 PM, are not significantly different between disease groups and present a major morning peak and a second lower afternoon peak, as represented by a density plot. E: An initial global gene expression analysis was carried out regardless of the pathological state of the liver, by comparing samples collected in the morning (AM) or in the afternoon (PM). A Volcano plot is shown representing gene expression fold-changes (AM vs. PM) and corresponding p-values obtained using DEseq2 considering sex as a confounding factor. **F:** KEGG pathway enrichment was calculated for the 1,660 genes whose expression was significantly different (adjusted p-value<0.05) between AM and PM. **E,** F:fontsizewasadjusted forclarity purpose.

## RESULTS

### Time-of-day is a major factor affecting gene expression in human liver

A human liver biopsy collection from patients with obesity undergoing bariatric surgery (HUL cohort, NCT 01129297), for which the exact daytime of the liver biopsy was studied (Figure 1A). Key clinical parameters of the 290 patients are summarized in Table 1. Based on histological parameters of liver biopsies (steatosis, hepatocyte ballooning, lobular inflammation), patients were grouped into NAFLD stages (labelled “HN”, histologically normal; “NAFL”, isolated hepatic steatosis; or “NASH”, steatosis, lobular inflammation and ballooning) following the indicated decision tree (Figure 1B). The proportion of men in this cohort of morbidly-obese diabetic patients increased with NAFLD severity, rising from 16% (HN) to 38% (NASH) with an average CRN NAS score rising from 0 to 5, respectively. Since sex is an important bias (*18*), this was considered during further analysis (see below). Patients in the NAFL and NASH groups were slightly older than those in the HN group, and expectedly had also higher insulin resistance on average.

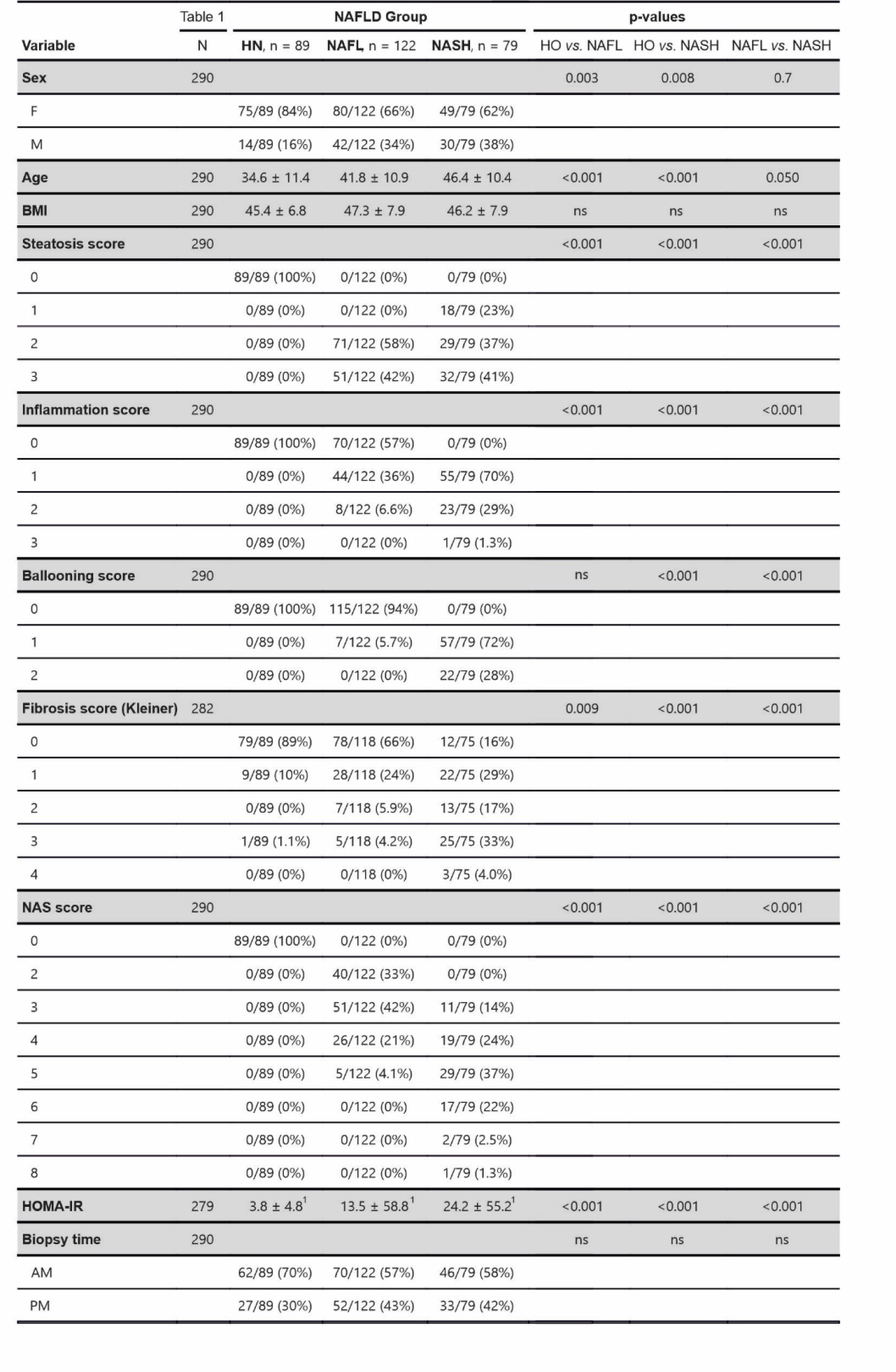
Biometric and biochemical parameters of the HUL sub cohort. The main biometric, biochemical and liver histological features of selected patients are indicated. F: women, M: men; BMI: body mass index; NAS: NAFLD Activity; HOMA-IR: Homeostatic Model Assessment for Insulin Resistance; AM: ante meridiem, PM: post meridiem. Continuous values are expressed as mean ±SD. Inter-group comparisons were performed using the unpaired Wilcoxon test for continuous variables (age, BMI, HOMA-IR) and Fisher’s exact test forthe remaining categorical variables.

The source of variation in gene expression levels was estimated by a multivariate analysis of variance (ANOVA) (Figure 1C). The F-ratio (ratio of the between-group variance to the within group-variance) confirmed sex and group (*i.e.* NAFLD stage) as the main sources of variation as previously reported (*18*), but very interestingly identified biopsy time (AM *vs.* PM) as the third most significant source of variation (Figure 1C), calling for a more in-depth analysis of time-dependent transcriptomes and their evolution during NAFLD progression. Age had only a minor contribution to signal variation. The distribution of available biopsy times (Figure 1D) revealed a daytime window of about 8 hours and indicated that biopsies were predominantly (>60%) taken in the morning with a first peak around 9:30 AM and a second, lower peak around 2:30 PM, due to the logistical schedule of surgical interventions. There was, however, no significant difference in biopsy time distribution between the groups (Table 1). Since preliminary tests excluding biopsies collected between 11am and 1pm did not modify the outcomes of statistical analysis, the “AM“ subgroup was defined as biopsies taken before noon (12:00 AM), the “PM” subgroup as biopsies taken after noon.

Gene expression profiles were thus compared by RNA-sequencing of morning (AM) *vs.* afternoon (PM) samples. Differentially expressed genes were identified using DEseq2 independent of the NAFLD status, but correcting for sex. Transcript counts for 1,660 genes were significantly different (Benjamini-Hochberg adjusted p-value <0.05; 4,375 genes before adjustment) between AM or PM biopsies (Figure 1E). Among the 100 top hits were most of the CCGs (*PER3, ARNTL/BMAL1, NPAS2, NR1D1, NR1D2, PER2, CRY1, PER1*), clock-related genes (*CIART*, *DBP, NFIL3)* (Figure 1E) as well as circadian-regulated genes involved in lipid metabolism (*PPARD*, *LIPG, LPIN2…)*. Because patients were fasting from midnight irrespective of surgery time, genes implicated in hepatic gluconeogenesis (*G6PC, PCK1, SGK2…)* were, as expected, higher expressed in afternoon samples (Figure 1E and Supplementary Table 1). In contrast to nocturnal rodents, genes from the negative limb of the clock displayed lower expression in the afternoon (*PER3, NR1D1, NR1D2, CIART…*), whereas genes from the positive clock limb were higher expressed in the afternoon (*ARNTL/BMAL1, NPAS2…*)(Figure 1E). Globally, genes displaying AM *vs.* PM differences were significantly enriched for the KEGG pathways “circadian rhythm”, “PPAR signaling pathway”, carbohydrate and lipid metabolic pathways, as well as cellular architecture and communication, among others (Figure 1F).

Taken together, our data show that an 8-hour time frame allowed the detection of significant changes in time-of-day-dependent liver gene expression, with a large proportion of varying transcripts functionally related to circadian rhythmicity.

### Time-dependent genes vary between NAFLD stages

We next examined whether time-of-day-dependent distributions of gene expression would differ between the histological states “HN”, “NAFL” and “NASH”. In order to achieve statistical power and obtain robust and exhaustive lists of time-dependent gene expression over the available daytime window for each stage, we used 3 complementary statistical methods analyzing different aspects of gene expression distribution (differential expression, partial Spearman correlation, Kolmogorov-Smirnov test), the results of which were agglomerated by a Fisher test to yield a combined p-value for each gene (Figure 2A-C and Supp. Figures 1 and 2). The 3 types of analyses are graphically exemplified for the core clock gene *ARNTL/BMAL1* (Figure 2A-C) which served as positive control to validate our approach, as it is among the most highly time-dependent genes regardless of the histological group. First DEseq2 (Figure 2A), which relies on a negative binomial distribution of gene expression, was used to identify differential expression between 2 conditions (AM *vs.* PM as in Figure 1). Second, partial Spearman correlation was computed between *ARNTL* gene expression and daytime (Figure 2B). Both approaches integrated sex as a confounding factor. Third, the Kolmogorov-Smirnov test was employed to determine whether AM and PM *ARNTL* expression distribution followed a similar law and are thus similar in shape (Figure 2C). Finally, the Fisher combined probability test or “Fisher’s method” was used as a meta-analysis method for p-value combination: individual raw p-values resulting from each statistical test were agglomerated into a single p-value per group (Figure 2D). The detected expression profile of *ARNTL*, of other CCGs (Supp. Figure 1) and of all other transcripts (Supp. Figure 2), clearly confirmed that the available time window was sufficient for robust time-of-day analysis of gene expression. After adjustment for multiple testing (FDR <0.01), a total of 2,738 genes were found to be time-dependent. When considering genes with an absolute fold change greater than 1.2 (AM *vs.* PM), this number was reduced to 1,427 (Figure 2E). The vast majority of these time-dependent genes (TDGs) were strikingly distinct when comparing the 3 patient groups, and only less than 10% (132 genes) were common to all 3 groups (“common TDGs”) (Figure 2E). However, most CCGs (*ARNTL, NR1D1/2, NPAS2, CRY1, PER1/2/3, DBP, CIART*) were found in this common intersection (Supplementary Table 1). TDG repartition outside of this core set was strongly unequal between groups, with ≈50% (558) of non-shared TDGs found in HN, ≈35% (392) in NAFL and less than 15% (177) in NASH livers. Along the same line, we found that AM to PM fold changes of common TDGs were on average decreased in NAFL and even more in NASH compared to HN (Figure 2F). These differences are illustrated for a selection of TDGs with AM- PM differences decreasing in NASH (*PCAT18, ARMC4, SIK1B*) and TDGs with AM-PM differences increasing in NASH (*CYP4Z1, RHOBTB1, PPIAP71*)(Figure 3A,B).

**Figure 2:**
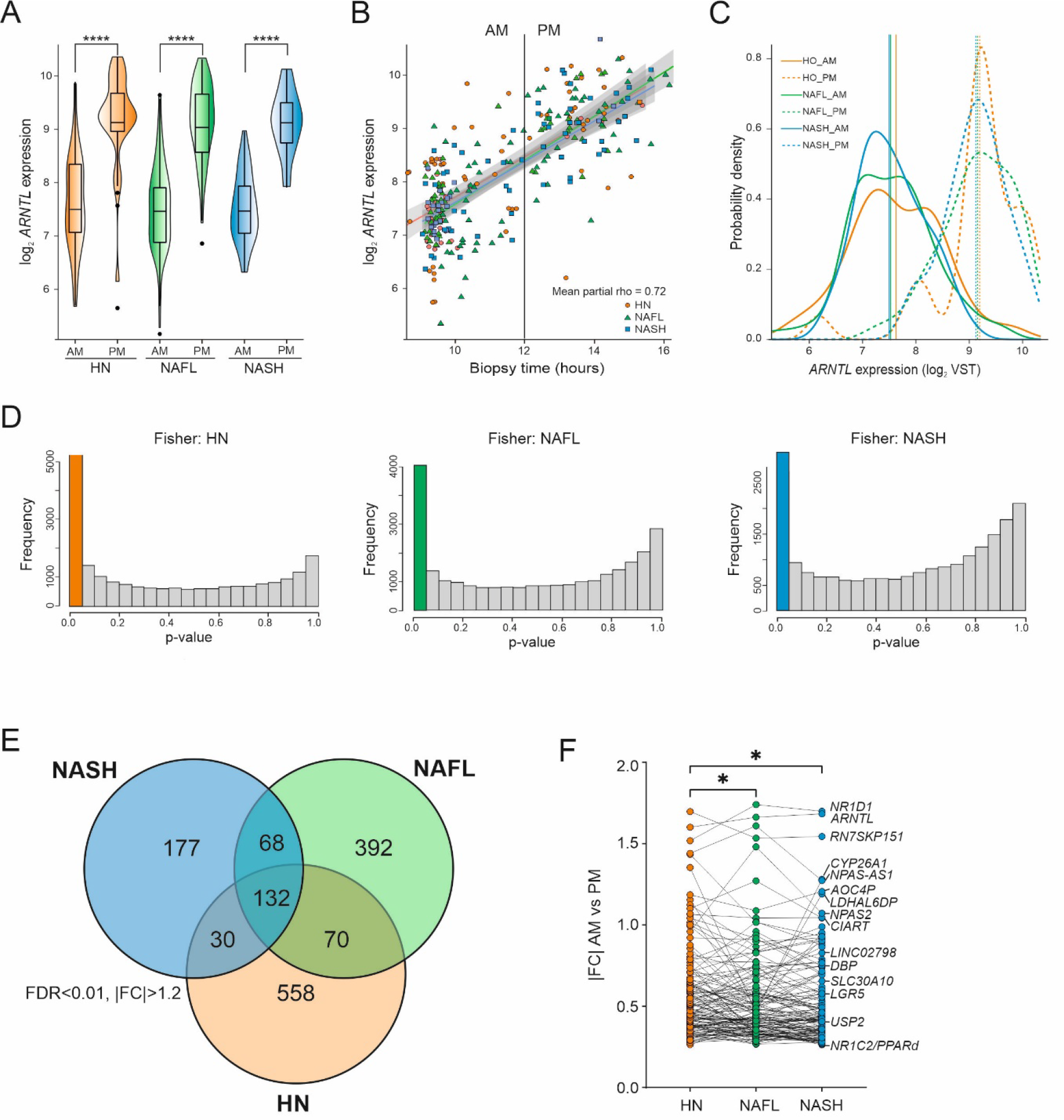
**A multi-test method to identify time-of-day-dependent genes (TDGs) within a restricted time window. A­C:** A combination of three different statistical methods was used to detect differences in gene expression depending on daytime of sample collection, illustrated here with the core clock gene ARWTÍ as an example; standard ana lysis with DEseq2 to compare AM vs. PM and correcting for sex as confounding factor **(A,** violin plot), partial Spearman correlation between gene expression and biopsy daytime **(B,** dot plot with linear tendency lines) and Kolmogorov- Smirnov test for comparison of AM or PM gene expression distributions **(C,** density plot). **D:** Fisher’s agglomeration method: in order to obtain a single p-value from combining the three abovementioned statistical tests, Fisher’s agglomeration method was applied to each gene within each of the 3 liver disease groups. Frequency histograms of uncorrected Fisher p-values are shown for each group and the first (colored) bar of each group histogram indicates genes with p<0.05. **E:** After applying p-value correction for multiple testing and stringent filtering of genes with a Fisher FDR<O.Ol and absolute AM-PM fold-changes greater than 1.2, 1,427 robustly time-dependent genes (TDGs) were identified. These are distributed unequally between liver disease groups as shown by a Venn diagram. **F:** The 132 TDGs overlapping between liver disease groups (“common TDGs”, central intersection in panel C) contained most core-clock genes and had overall reduced absolute AM-PM fold-changes (* indicates p<0.05 using ANOVA and Fisher’s LSD post-hoc test). VST: variance-stabilizing transformation.

**Figure 3:**
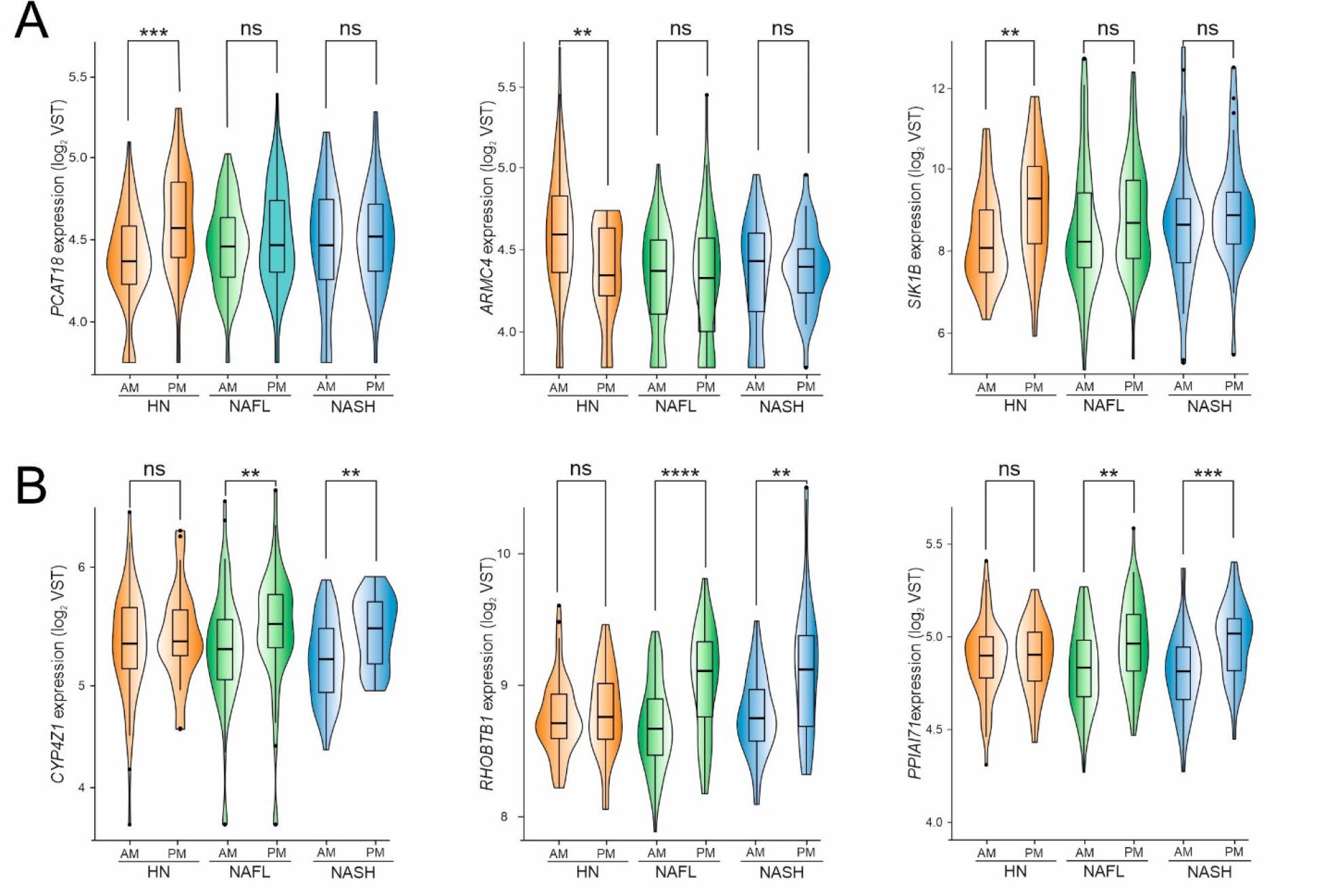
**Example distributions of time-dependent genes (TDGs) altered in NASH.** Certain TDGs exhibit greater AM- PM fold-changes of gene expression in healthy livers (HN) compared to NASH *i.e.* they lose their time-dependency in NASH, as shown in panel **A.** Others have greater AM-PM fold-changes of gene expression in NASH and are thus becoming time-dependent in NASH, as shown in panel **B.** Statistical tests were Kruskal-Wallis tests followed by unpaired Wilcoxon post-hoc test for AM-PIVI comparisons in each group (*p<0.05, **p<0.01, ***p<0.005, **♦*, p<O.OOl).

Enrichment analysis for KEGG pathways showed that common TDGs, *i.e.* identified in all three groups, were strongly enriched for terms like “circadian rhythms” as expected from the content in transcripts coding for CCGs (*NR1D1*, *NR1D2*, *ANRTL*, *CIART*…), but also for metabolic regulatory pathways like the PPAR and FoxO pathways (Figure 4, Supp. Figure 3), illustrated by genes such as *S1PR1, G6PC1, PCK1, SGK2, FASN, AQP7* and *PPARD*. TDGs identified as unique to the HN group were also enriched, albeit to a lesser extent, for pathways linked to circadian rhythm (notably including *CLOCK*) as well as to fatty acid and amino acids metabolism, but the most highly represented pathway was “gap junctions” (Figure 4, Supp. Figure 3). This pathway was characterized by genes such as *PDGFB, MAP2K1* and transcripts encoding for tubulins *TUBA1C/8, TUBB, TUBB1/2B* (Supp. Table 1), suggesting that epithelial barrier integrity/permeability homeostasis, which is known to be disturbed in NAFLD (*30*), requires an oscillating expression of these genes in healthy conditions. Genes unique to the NAFL group (Supp. Table 1) were mostly linked to metabolism of lipids and fatty acids (*DGKG, PLA2G4B/5, LPIN2/3, ETNK2, ETNPPL, PLPP4, FADS1/2, CYP2C8, GDPD1, SCD*) and also to metabolism of peptides and amino acids (*DNMT3B, GCLM, SDS, PSAT1, GNMT, ALDOC, SDS, GPT2, CSAD, UPB1*)(Figure 4, Supp. Figure 3). Lastly, TDGs specific to the NASH group (Supp. Table 1) were highly enriched for signaling by calcium, cAMP or neurotransmitters (*ADRB2, DRD1, GRM1, NTRK1, NTSR1, P2RX7, RYR2, TBXA2R, CACNA1H, SSTR5, TBXA2R, FFAR2, SUCNR1)* as well as for lipolysis (*ADRB2, IRS1, PNPLA2*)(Figure 4, Supp. Figure 3). Taken together, these data show that the temporal pattern of gene expression in homeostatic conditions is strongly affected by the disease state and indicative of compromised cellular communication and metabolic pathways.

**Figure 4:**
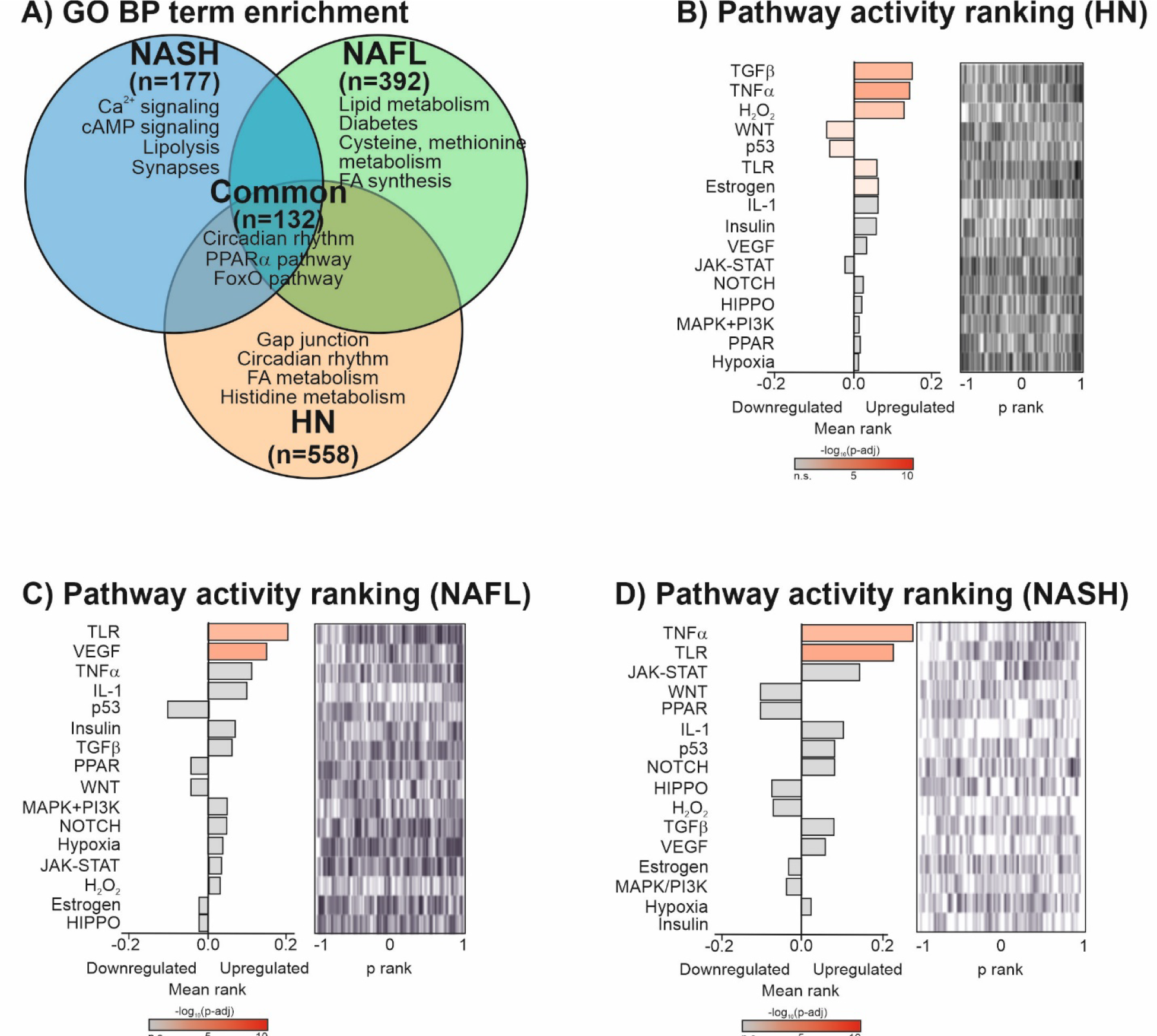
Term enrichment analysis of time-dependent genes (TDGs) and upstream regulatory pathway prediction. A: Biological term enrichment analysis. TDGs identified in Figure 2 were analyzed for enrichment of gene ontology (GO) terms related to biological processes (BP) using the online gene enrichment analysis platform Metascape. Significantly enriched GO-BP clusters were manually collapsed for visualization purpose and top hits are indicated. B-D: Pathway activity ranking. The Speed2 tool was used to quantify whetherTDGs are enriched for pathway-deregulated signature genes. Each pathway is represented as a bar showing the mean rank of the query list. The “bar code” plot shows the distribution of genes from the query list in the ranked reference signatures.

Inferring upstream regulatory cues or altered biological processes may be achieved by comparing differentially expressed gene lists, identified here by our combinatorial approach, to annotated gene sets (Figure 4A) or to consensus gene expression patterns induced by a given perturbagen (Figure 4B-D). TDGs specific to each stage (HN, NAFL, NASH) were first analyzed for term enrichment from the Gene Ontology Biological Pathways (GO BP) database (Figure 4A). Common TDGs displayed, as expected, significant enrichment for circadian- and metabolic-related processes. TDGs specific for normal livers (HN) emphasized tissue functions centered around cell to cell communication (gap junctions) and metabolism, whereas fatty livers (NAFL) displayed increased involvement in lipid, glucose and amino acid metabolism. TDGs specific for NASH were significantly associated to intracellular signaling (Ca^2+^ and cAMP).

Speed2 (Signaling Pathway Enrichment using Experimental Datasets (*31*)) analysis allows probing gene lists against ranked gene signatures for 16 signaling pathways, with the aim of identifying upstream signaling mediators. Ranked signatures suggested that cues in homeostatic (HN) conditions could be TGFβ, TNFα, oxidative stress, TLR and estrogen (Figure 4B). In NAFL and NASH conditions, this pattern shifted towards a more limited signaling pathway panel with similar statistical significance, which included either TLR and VEGF (NAFL) or TNFα and TLR (NASH)(Figure 4C and 4D respectively). Although causative links cannot be proven, this data could reflect a loss of physiological rhythmic function(s) in NAFL and NASH livers, which in turn gain rhythmic functions associated to pathogenic immune and proliferative stimuli and responses.

### NAFLD stages modify time-of-day changes in liver metabolites

The results of our gene expression analysis suggested that metabolic pathways are altered in a time-of-day-dependent manner as NAFLD progresses, with an enrichment in amino acid- and lipid metabolism-regulating genes (Supplemental Figure 3). Therefore, an unbiased tissue metabolomic study by LC-MS was performed on liver samples of the 290 patients included in the transcriptomic study. Similar to the gene expression analysis, a global approach was initially employed to evaluate overall time-of-day dependence of tissue metabolite levels regardless of the NAFLD status. This global analysis identified ≈220 metabolites whose amounts were significantly different in morning and afternoon biopsies (DEseq2 corrected for sex, FDR<0.1) (Figure 5A, Supplemental Table 2). Visual inspection of the volcano plot highlighted intermediates of lipid β- oxidation (carnitine derivatives), amino acids (kynurenate, oxo-arginine…) as being differentially detected in AM *vs.* PM livers (Figure 5A). It also confirmed the marked fasting status of patients undergoing surgery in the afternoon by an increased hepatic content in 3-hydroxybutyrate (BHBA). A biological term enrichment of identified metabolites confirmed that the majority of the identified metabolites belonged to amino acid and lipid and fatty acid metabolic pathways (Figure 5B).

**Figure 5:**
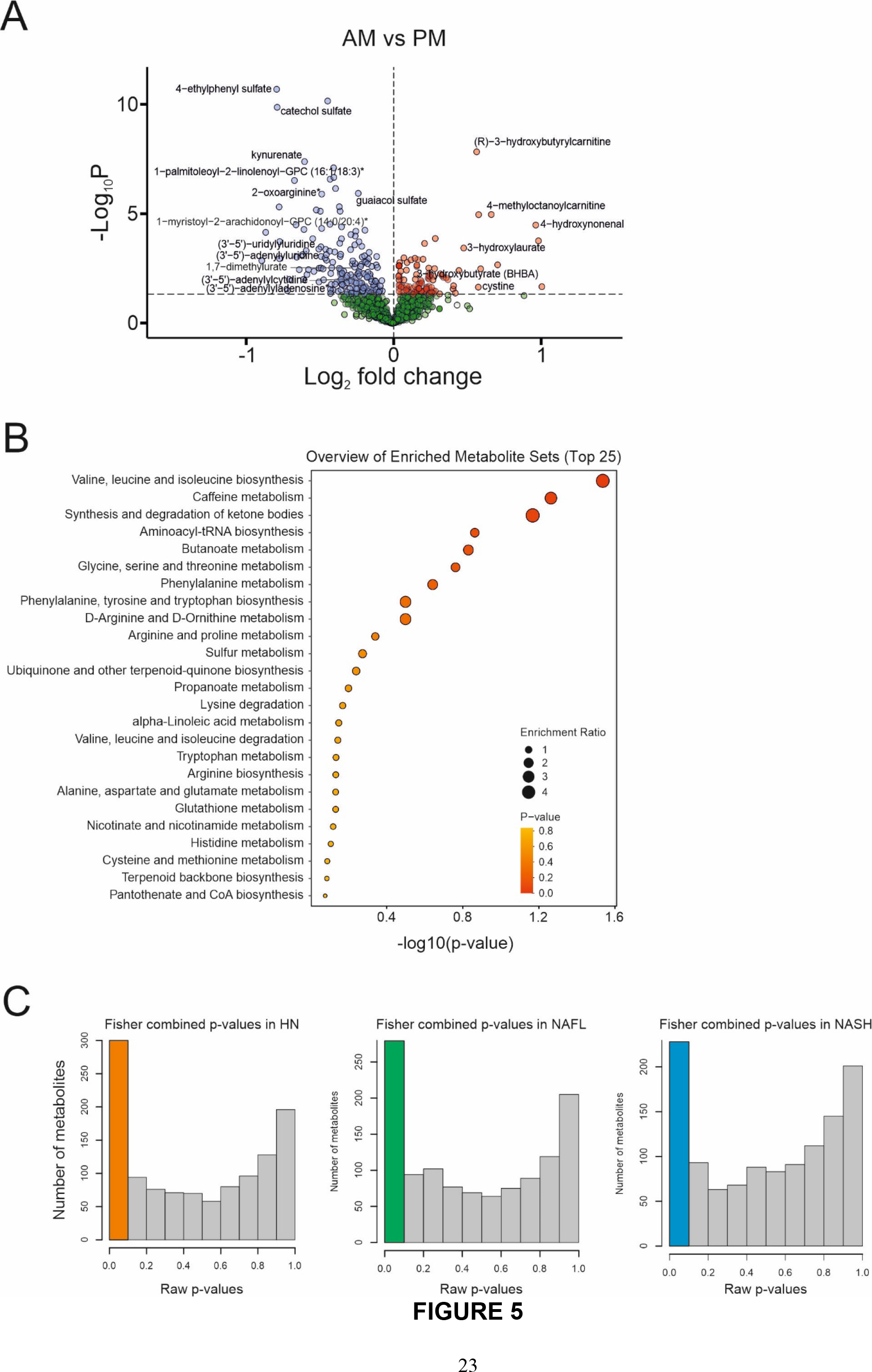
**Identification of time-dependent liver metabolites measured by LC-MS. A:** Similar to gene expression analysis (see Figure 1), an initial global analysis was carried out to compare abundances of liver tissue metabolite in biopsies taken in the morning (AM) or afternoon (PM), regardless of the disease state of the liver. A Volcano plot is shown representing AM-PM fold- changes of metabolite abundances and corresponding p-values obtained using DEseq2 considering sex as confounding factor. **B:** Enrichment of KEGG metabolic pathways was calculated for the 220 metabolites whose abundance was significantly different (DEseq2 adjusted p-value <O.l) between AM and PM. **C:** As for gene expression analysis (see Figure 2), liver biopsies were divided into three groups according to liver disease state: “HN” (histologically normal), “NAFL” (benign steatosis only) or “NASH” (steatosis + inflammation). A combination of three separate statistical tests (DEseq2, partial Spearman correlation, Kolmogorov-Smirnov) with p-value agglomeration by Fisher’s method was used to identify time-dependent metabolites (TDMs) within each group. The resulting uncorrected Fisher p-values are shown as frequency histograms and the first (colored) bar of each group histogram indicates metabolites with p<O.l.

To highlight a possible time-of-day differential representation of metabolites between NAFLD groups, we again combined the 3 statistical approaches as described for gene expression analysis (DEseq2, Spearman correlation, Kolmogorov-Smirnov test) followed by Fisher’s agglomeration for a robust identification of time-dependent metabolites (TDMs) (Figure 5C). A total of 251 TDMs were identified using this method (combined FDR<0.1), out of which only 14 (6%) were common to all 3 NAFLD groups (Figure 6A, 6B). These common TDMs included amino acids such as proline and threonine, several fatty acids and the ketone body component β- hydroxybutyrate (Figure 6B and Supp. Figure 4A-F). KEGG metabolic pathway enrichment analysis revealed that these common TDMs were most significantly associated to metabolism of amino acids (arginine, proline, threonine)(Figure 6B and Supp. Figure 4A-C). Synthesis of ketone bodies was also identified (Figure 6B), in agreement with the identification of the PPARα pathway based on gene expression patterns (Figure 4) and the detection of 3-hydroxybutyrate (Supp. Figure 4D).

**Figure 6:**
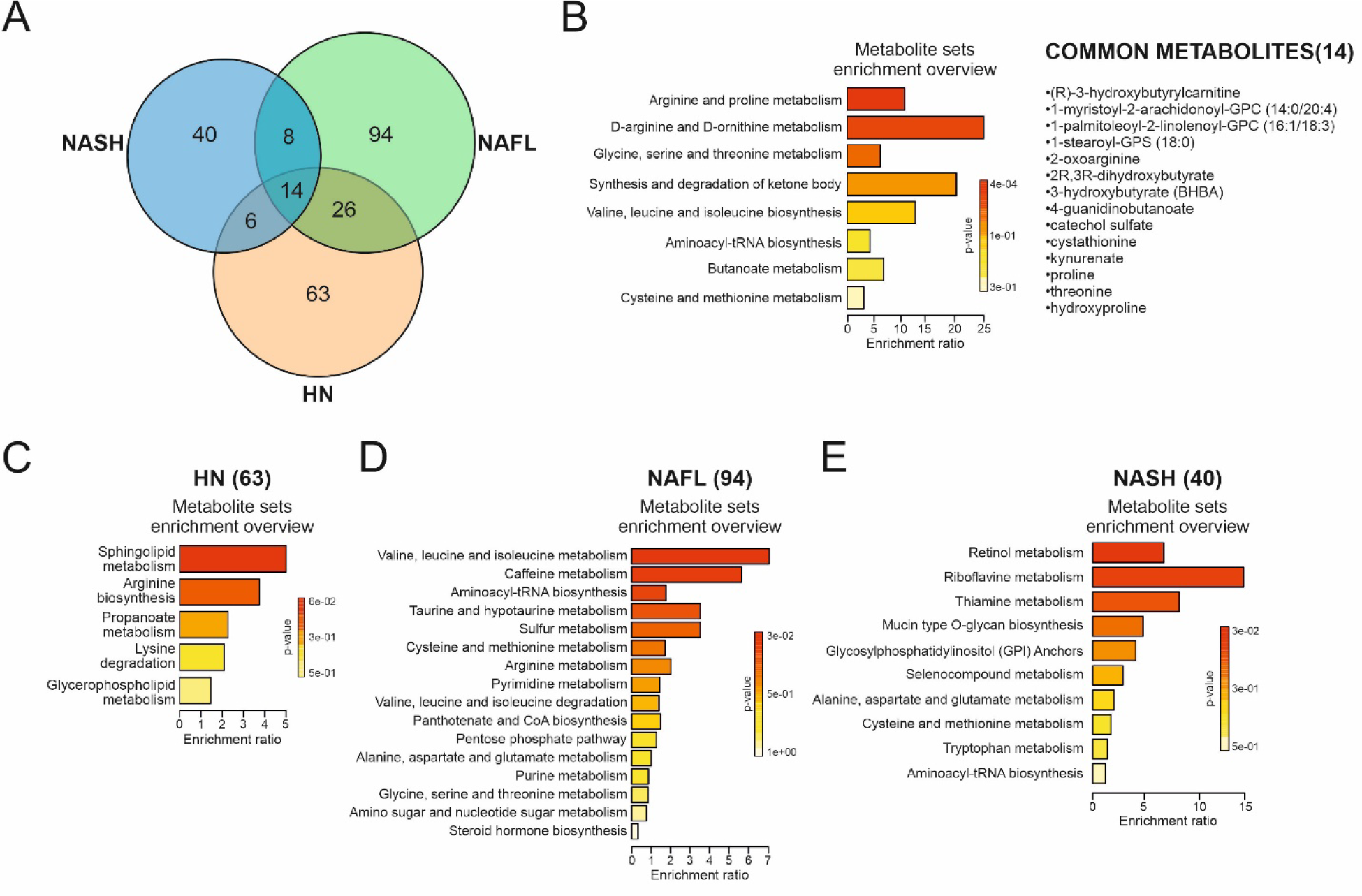
Distribution of time-dependent liver metabolites (TDMs) among liverdiseasegroupsand KEGG metabolic pathway enrichment. After correction for multiple testing, metabolites with a Fisher FDR<O.l were considered robustly time-dependent (TDMs). The Venn diagram shows the distribution of these 23S TDMs among liver disease groups, highlighting major differences between groups, KEGG metabolic pathway enrichment of common (central intersection) or group-specific TDMs was calculated usingthe online metabolomics analysis platform MetaboAnalyst.

Among the 197 stage-specific TDMs, 31% were specific to HN, 47% to NAFL and 20% to NASH (Figure 6A and Supp. Table 2). TDMs specific to HN livers, and thus lost at the NAFL and NASH stages, were mainly associated to metabolism of sphingolipids (Figure 6C and Supp. Table 2) such as CDP-choline and sphinganine (Supp. Figure 4G, H). Glycerophosphoethanolamines, glycerophosphocholines as well as cholesterol, amino acid and pyrimidine derivatives were also identified as time-dependent in normal livers (Supp. Table 2 and Supp. Figure 4I-L).

A similar enrichment analysis identified amino acid metabolic pathways (branched-chain, sulfur-containing, arginine, taurine) (Figure 6D, Supp. Figure 5A, B and Supp. Table 2) as time-dependent in NAFL livers. Visual inspection of NAFL TDMs also identified a carnitine precursor (N6,N6,N6 trimethyl-lysine, Supp Figure 5C) and derivatives (Supp. Table 2 and Supp. Figure 5D, E) which could reflect an altered fatty acid oxidation activity. Finally, NASH-specific TDMs were enriched mainly for vitamin, glycan and glycosylphosphatidylinositol (GPI) metabolic intermediates (Figure 6E, Supp. Figure 5 and Supp. Table 2).

Taken together, these analyses highlight the disruption during NAFLD progression of time-of-day-dependent bioactive phospholipid metabolism and of amino acid biotransformation pathways. Intriguingly, we also noted that established PPARγ ligands of the linoleic acid class [9,10 DiHOME (*32*) and 9- and 13-HODE (*33*)] displayed a differential abundance in AM *vs.* PM NAFL and NASH livers, with estimated concentrations in the 10-100µM range probably sufficient to activate PPARγ (Supp. Figure 5F, J).

### Integrative analysis of time-dependent genes and metabolites

We next performed an integrative analysis of the transcriptomic and metabolomic data at the pathway level using the KEGG database. As a first step, this analysis combined TDGs and TDMs specific to either HN or NASH stages and common TDGs and TDMs, irrespective of their relative time-of-day direction of change in order to identify associated transcriptomic and metabolomic conditions operating in normal and NASH livers (Figure 7A). TDGs and TDMs characterizing the HN stage were enriched for metabolic pathways related to lipid and amino acid metabolism, while most of them were not detected at the NASH stage, or with a decreased significance (Arg and Pro metabolism, glycerophospholipid metabolism). Linoleic metabolism was associated to the NASH stage (Figure 7B, C). HN-or NASH-enriched pathways (glycerophospholipid and linoleic pathways, respectively) were further detailed for daytime variation of associated TDGs and TDMs. The glycerophospholipid pathway was characterized by an increased abundance in AM livers of 3 out of 4 detected diacylglycerol (DAG) species specifically at the HN stage. Glycerophosphocholine (GPC) intermediates (CDP-choline, GPC) displayed stage-specific time-of-day variations that were not correlated to the occurrence of glycerophospholipid species (X-GPC) (Figure 7D). Higher DAG species abundance in the morning did not correlate with HN TDG expression changes in transcripts encoding enzymes involved in this metabolic pathway, with the exception of the patatin-like phospholipase domain–containing protein 3-encoding gene (*PNPLA3*). PNPLA3/adiponutrin has acyltransferase activity, increasing the formation of phosphatidic acid (PA) from lysophosphatidic acid (LPA), which may lead to more DAG synthesis. It might also reflect PNPLA3 hydrolytic activity on triacylglycerol molecules, favoring DAG accumulation. Along the same line, we compiled TDG and TDM data related to linoleic acid metabolism (Figure 7E). The increased abundance of the PPARγ ligands 9- and 13-HODE in the afternoon at the NASH stage was mirrored by the gene expression of *ALOX15*, a dioxygenase catalyzing the synthesis of these 2 hydroxyoctadecadienoic acids which was increased in the morning. A similar lack of correlation was observed between 9,10-DiHOME hepatic content and the expression of the linoleic acid-converting *CYP2C8* at the NAFL stage.

**Figure 7:**
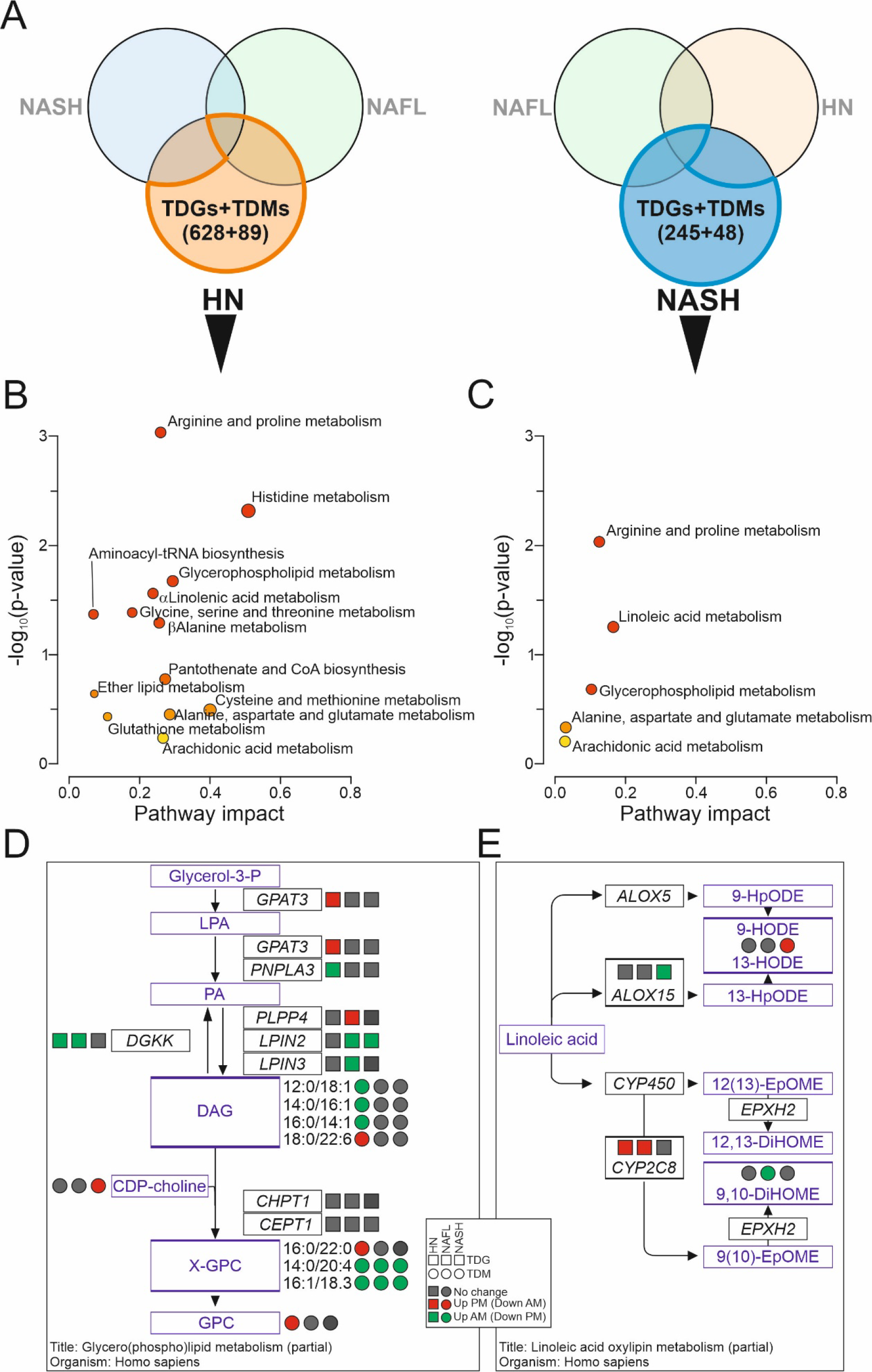
Integrative analysis of time-dependent genes (TDGs) and metabolites (TDMs) and alterations in NAFLD. Common (central intersection) combined to the HN or the NASH group-specific TDGs and TDMs (A) were analyzed associatively for enrichment of KEGG metabolic pathways using the online metabolomics analysis platform MetaboAnalyst, revealing links between metabolic pathways as defined by time- dependent gene expression patterns and metabolomic profiling (B, C). Reconstitution (partial) of metabolic pathways displaying time of the day-dependency in the HN (D) or NASH (E) group. LPA: lysophosphati die acid, PA: phosphatidic acid, DAG: diacylglycer¤l, CDP-choline: cytidine-diphosphate-choline, PC: phosphatidylcholine, GPC glycerophosphatidylcholine, X-GPC: acyl-conjugated GPC, 9-HpODE: 9- hydroperoxy octadecadienoic acid, 13-HpODE: 9-hydroperoxy octadecadienoic acid, 9-HODE: 9- Hydroxyoctadecadienoic acid, 13-HODE: 13-Hydroxyoctadecadienoic acid, 12(13)-EpOME: 12(13)-epoxy-9Z- octadecenoic acid, 9,10-EpOME: 9(10)-epoxy-12Z-octadecenoic acid, 12,13-DİHOME: 12,13-dihydroxy-9- octadecenoicacid, 9,10-DiHŨME: 9,10-dihydroxy-9-octadecenoicacid.

These results suggest that time-dependent metabolite variation in these pathways is delayed with respect to gene regulation, and/or controlled by post-transcriptional processes. Still, these observations strongly confirm the presence of time-of day variation in hepatic gene expression and metabolome, and identify several new oscillating metabolites at the NASH stage (Figure 7 and Supp. Table 2).

## DISCUSSION

Ideal conditions for chronobiological studies such as carried out on rodents imply obtaining multiple replicates at 2h-intervals over a total period of 24 or even 48 hours, within a controlled environment including exposure to light and food. These conditions are hardly achievable when studying humans, which is the main reason why circadian rhythmicity remains poorly analyzed in humans, and particularly in internal organs (*34*). While human circadian rhythms are well appreciated by GWAS studies and the phenotypic manifestation of disturbed cyclic processes such as sleep, light exposure and eating patterns (*35*), their study in healthy or pathological conditions is indeed hindered by ethical and technical constraints. Although most of the reported studies were carried out in a controlled experimental setup with regard to sleep behavior and food intake, samples were essentially collected serially at a limited number of time points and from a very small number of mostly healthy individuals. This is problematic given the high inter-sample variability of human biopsies which is even further amplified in the context of chronobiology.

In the current study, we reveal the first ever time-of-day-resolved human liver transcriptome with associated liver tissue metabolites using a 290-patient cohort. Although the available time window of liver biopsies was only about 8 hours, this temporal window was sufficient to robustly identify genes and metabolites varying over time (TDG and TDM respectively). We then went further in our analysis to highlight differences in the time-dependence of genes and metabolites between histological NAFLD groups defined by the presence of steatosis alone (“NAFL”) or combined with ballooning and lobular inflammation (“NASH”) and compared them to livers from histologically normal (“HN”) individuals. While a strikingly small fraction of genes and metabolites were identified as time-dependent in all three patient groups, these TDGs/TDMs were unequally distributed among the 3 groups. This indicates that while the expression pattern of CCGs, which were mostly found in the common group, is rather robust even in pathological conditions, the alignment of rhythmic biological processes such as intercellular communication (gap junctions) and metabolic regulations is disrupted upon NAFLD progression. However, none of the detected TDG in NASH patients belonged to the human and mouse core set of NASH/fibrosis-associated genes (*20*). Interestingly, the number of stage-specific TDGs (Figure 2E) decreases from the HN to the NASH stage, and concomitantly enriched pathways lessen (Supp. Figure 3), hinting at a loss of functional adaptability/(metabolic) flexibility. In high fat diet (HFD)-fed mice, transcripts gaining rhythmicity when compared to chow diet-fed mice are strongly enriched for glycerophospholid metabolism (*22*), similarly to NAFL patients (Supp. Figure 3), indicating convergent mechanisms for liver adaptation to dietary imbalance as often occurring in NAFLD. At the cellular level, Ca^2+^ fluxes are submitted to ultradian variations and coupled to metabolic regulations (*36*), and mishandled intracellular Ca^2+^ stores in NASH can significantly impact parenchymal and non-parenchymal liver cellular functions (*37*). A number of genes encoding for Ca^2+^ channel components or involved in intracellular Ca^2+^ signaling (*PKD1L1*, *TRPC1 P2RX7*, *FFAR2*, *ADRB2*, *CACNA1H*, *GRM1*, *NTRK1*) exhibited differential AM *vs.* PM expression specifically in NASH livers. Thus, our data show that in addition to the lipid-induced dysfunctional ER Ca^2+^ transport (*38*), *de novo* oscillation of the calcium handling process accompanies progression to NASH. *CLOCK* gene deletion affects metabolite oscillations in mouse liver (*25*) and our analysis showed a time-of-day differential expression of *CLOCK* exclusively in HN livers, which display a more diverse metabolic activity than NASH livers (Figure 7B). While hinting at a possible role of this component of the positive arm of the molecular clock, the examination of individual metabolites nevertheless showed little overlap between mouse *CLOCK*-dependent metabolites and HN-specific metabolites. This lack of exact concordance can be explained by species-specific mechanisms, distinct effects of gene deletion *vs.* loss of time-dependent expression and/or technical bias.

While examining the coherence between NAFLD state-specific liver transcriptomes and metabolomes, we observed little correlation between enzyme-encoding genes and metabolite abundance. This disconnection is not unprecedented and was also observed in a highly standardized mouse study which minimizes the variability typically observed in human samples (*22*). The narrow time window of our study may explain in part this lack of correlation as transcripts are likely to precede metabolite production, hence affecting our statistical approach for the PM sub-cohort. It may also indicate significant time-of-day-dependent translational control (*39*) as well as post-translational modifications regulating enzyme activity, both processes which are not captured by our analysis.

Finally, another point of convergence between our and mouse studies is the differential abundance of linoleic acid derivatives and PPARγ ligands 9,10 DiHOME, 9-HODE and 13-HODE in NAFL and NASH livers, respectively. The PPARγ-encoding gene *NR1C3/PPARG* itself did not display significant oscillatory expression in human liver, contrasting with HFD mouse livers (*22*). Whether estimated intracellular concentrations of these compounds are indeed sufficient to differentially activate human liver PPARγ and have a causative role in hepatic transcriptional reprograming requires further in-depth investigation.

On the one hand, it is remarkable that many of the biological processes and pathways shown to be affected by NAFLD in other studies, identified on the basis of changes in gene expression or metabolite abundance levels regardless of time, also display altered time-dependent expression profiles in our study. On the other hand, we identified many novel potential links between genes with deregulated timed expression and NAFLD pathogenesis, which were not previously considered by standard analyses which did not integrate time-of-day as a variation factor. It is probably the combination of both types of deregulations that underlies the deeply disturbed liver functions once NASH is declared. Conversely, some of the changes detected when time-of-day information is absent or ignored may turn out to be artefacts, as time-of-day as a biological variable might not be equilibrated between groups.

Considering that the time-of-day dependence of transcript/protein/metabolite measurements was previously neglected or ignored in nearly all human NAFLD studies, our findings here reveal a significant impact of time-of-day on many relevant pathogenic processes. As such, differences found between sample groups in cohort studies might reflect a previously under-appreciated bias in sampling time between groups. Further investigation is needed in this regard, particularly when studying human material. In any case, a new generality should be that sampling daytimes (or *Zeitgeber* times) must be carefully recorded and included in post-hoc analyses whenever possible.

### Strengths and limitations of the study

This study has both strengths and limitations. This first-of-its-kind study revealed time-of-day transcriptomic and metabolomic alterations in human livers as a function of the histologically-proven NAFLD stage. It uses a large cohort allowing both the selection of biopsies to adequately encompass control, NAFL and NASH cases and robust statistical analysis. There is however a number of limitations, inherent to the observational nature of the study. The narrow time window for biopsies collection (8 hours) precludes the assessment of a 24h diurnal rhythmicity, hence of the integrity of the molecular clock. The unavoidable fasting duration could also be a confounding factor.

## MATERIALS AND METHODS

### Liver biopsies from the HUL cohort

The Hopital Universitaire de Lille (HUL) cohort, also known as the Biological Atlas of Severe Obesity (ABOS) cohort was established as from 2006 by the University Hospital of Lille, France (ClinicialTrials.gov: NCT01129297) from severely and morbidly obese patients visiting the Obesity Surgery Department. All patients of the cohort fulfilled criteria for, and were willing to undergo, bariatric weight-loss surgery (for details, see (*18*)). The protocol required that patients were fasting from midnight to surgery time. During the surgical procedure, wedge biopsies were taken from the liver to be immediately snap-frozen and the exact time of the biopsy was noted. A total of > 1,500 patients are currently included in the HUL cohort, amongst whom 319 were selected to build a sub-cohort with complete clinical, biometric parameters and a robust histological NAFLD classification of quality-controlled biopsies eliminating all intermediary NAFLD stages (see (*18*) and Figure 1B for more details). Both transcriptomes and metabolomes were obtained for these 319 patients with biopsy mass >100mg. Out of these, 290 had known biopsy time-of-day, ranging from 8am to 4pm, and were included in this study. Main clinical and histological characteristics are indicated in Table 1.

### Total RNA sequencing and data processing

Total RNA (800 ng) was used to perform library preparation using the KAPA RNA HyperPrep kit with RiboErase (HMR). Sequencing was done on the NovaSeq6000 system (Illumina) using a paired-end 2×75 bp protocol. Raw data were demultiplexed into FASTQ files using bcl2fastq v2.20.0.422. The quality of FASTQ files was checked using FastQC v0.11.9. An adapter trimming step was done using trimmomatic v0.39. Mapping of reads on the human genome (Hg38) was performed using STAR aligner (v2.7.3a). On average, 130 million reads accurately mapped against the human genome. Transcript counts were obtained using RSEM (v1.3.0).

### Liver metabolomics by LC-MS

All tissue samples were flash frozen and maintained at –80°C until processing. Sample preparation was carried out as described previously (*40*) at Metabolon, Inc. (Morrisville, NC, USA). Briefly, recovery standards were added prior to the first step in the extraction process for quality control purposes. To remove protein, dissociate small molecules bound to protein or trapped in the precipitated protein matrix, and to recover chemically diverse metabolites, proteins were precipitated with methanol under vigorous shaking, followed by centrifugation. The resulting extract was analyzed by four different LC-MS methods. Three types of controls were analyzed in concert with the experimental samples: samples generated from a pool of human plasma (extensively characterized by Metabolon, Inc.) served as technical replicate throughout the data set; extracted water samples served as process blanks; and a cocktail of standards spiked into every analyzed sample allowed instrument performance monitoring. Peak extraction and identification were automatically performed by comparison to Metabolon’s internal spectral library. Metabolite abundance was calculated from peak areas normalized to the internal standards. Data were scaled such that the median was equal to 1 and missing values were imputed with the minimum value.

### Bioinformatic analysis

All analyses were carried out using RStudio (v1.4.1106) with R (v4.1.0). Patient clinical data shown in Table 1 were analyzed using the package “gtsummary” (v1.5.0). For transcriptomic analysis, RSEM transcript counts of gene isoforms were rounded and summed per gene. Gene counts were prefiltered for low read counts to remove genes with row sums of read counts lower than the number of samples (thus with a mean read count per sample below 1). For differential expression comparing morning (AM, before 12:00am) versus afternoon (PM, after 12:00am) biopsies, the “DEseq2” package (v1.43.0) was used including also sex as a confounding factor in the design formula. To obtain robust lists of time-dependent genes, 2 additional statistical approaches were employed testing different aspects of the time dependence. Both approaches were applied after variance-stabilizing transformation (VST) of the gene count matrix from DESeq2. First, the Kolmogorov-Smirnov test (built-in “stat” package, v4.1.0) was employed to compare gene expression distribution between AM and PM. Second, partial Spearman correlation adjusting for sex (“ppcor” package, v1.1) was used to evaluate correlation between gene expression and time-of-day. The resulting 3 individual p-values from each approach were then agglomerated into a single p-value per NAFLD group for each gene, using Fisher’s method with the “metap” package (v1.7) and finally adjusted for multiple testing by applying a Benjamini-Hochberg correction (BH method, built-in “stats” package v4.1.0). Genes were considered as robustly time-dependent when the adjusted Fisher p-value (Fisher FDR) was lower than 0.01 with an optional cut-off on absolute fold-change AM *vs* PM above 1.2. The metabolomic analysis essentially followed the same procedure using processed/scaled abundance data as input and metabolites were considered time-dependent when the adjusted Fisher p-value was below 0.1 with no cut-off on AM-PM fold-change. Volcano plots were graphed with the “EnhancedVolcano” package (v1.12.0). Venn diagrams were made with the “VennDiagram” package (v1.7.1). All other plots were created with the “ggplot2” package (v3.3.5).

### Enrichment analysis

Time-dependent gene lists were analyzed for enrichment of Kyoto Encyclopedia of Genes and Genomes (KEGG) pathways and gene ontology (GO) biological process terms using Metascape 3.5 ((*41*), https://metascape.org/) with default settings. GO term clusters were manually renamed and/or regrouped together when judged pertinent and bubble charts were created using GraphPad Prism 9.3.0. Signaling pathway enrichment analysis was performed using the Speed2 online tool ((*31*), https://speed.sys-bio.net/) with the recommended standard Bates-test. Time-dependent metabolite lists were analyzed for enrichment of KEGG metabolic pathways using MetaboAnalyst 5.0 ((*42*), https://www.metaboanalyst.ca/) with default settings, as well as for joint-pathway analysis integrating metabolite and gene lists.

### Data visualization and illustrations

Graphs were generated as *.svg files using R packages mentioned above. Data were imported in CorelDraw2020 to assemble figures. Drawings in Figure 1A are from Renée Gordon, Victovoi, and Mikael Häggström, M.D. and were made available to the public domain via Wikimedia Commons with no restriction of use.

## Author contributions

Conceptualization: MJ, JTH, JV, FP, BS, PL; Software: MJ, JTH, JV, MD; Validation: MJ, JTH, JV, BD, VR, JE, BS, PL; Formal analysis: MJ, JTH, JV, JE, PL; Investigation: MJ, JTH, JV, BD, MD, AB; Resources: AB, PF, FP, BS, PL; Data Curation: MJ, JTH, JV, BD, MD, VR, PL; Writing: MJ, JTH, JV, BS, PL; Visualization: MJ, JTH, JV, PL; Supervision: BS, PL; Project administration: BS, PL ; Funding acquisition: FP, BS

## Data and materials availability

All data, code, and materials used in the analysis will be available in some form to any researcher upon reasonable request.

## List of Supplementary Materials

Supplementary Fig S1 to S5.

Supplementary Data files TableS1 and Table S2 (Excel files)

**Supplementary Figure SI.**
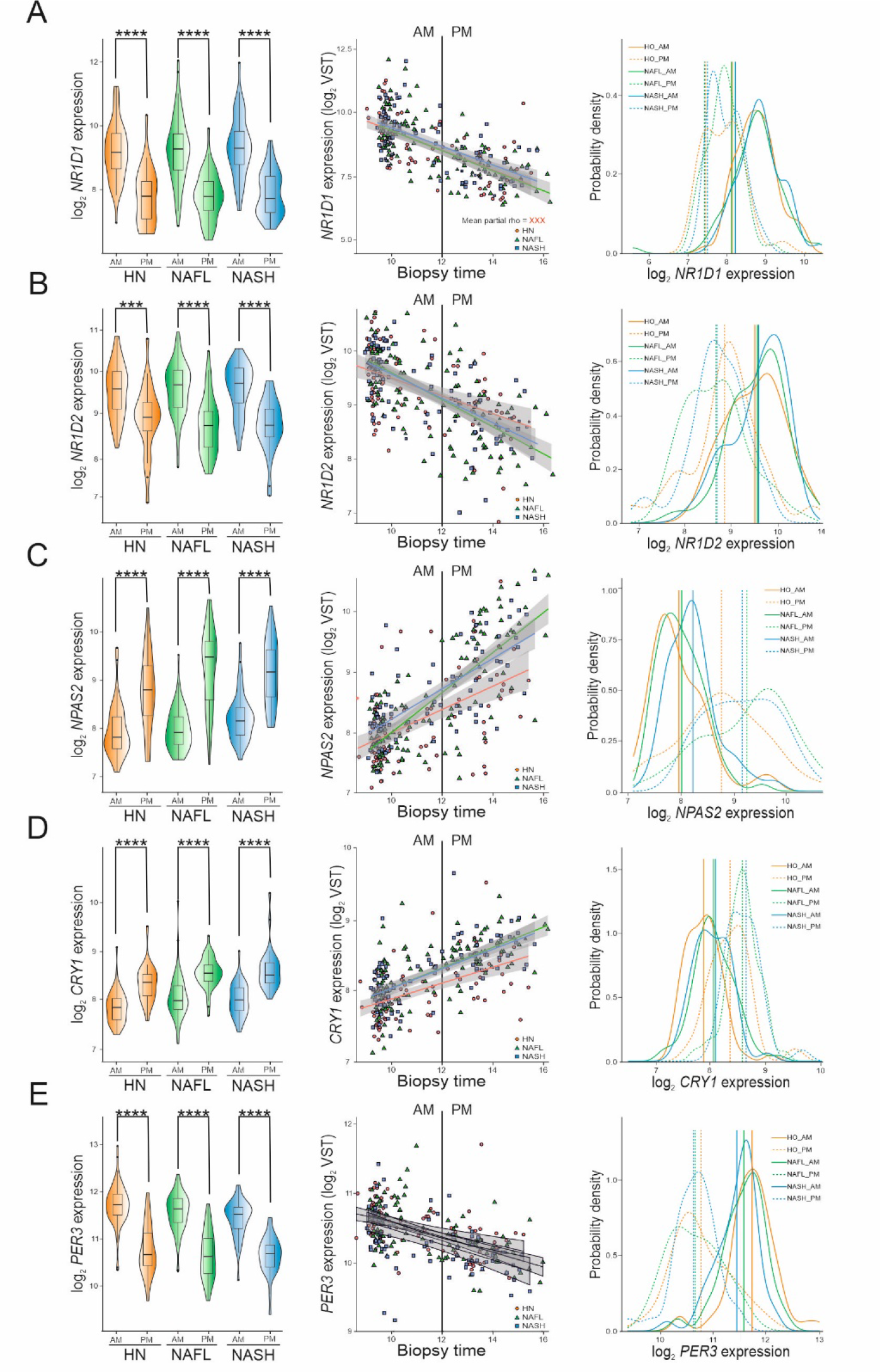
**Daytime-resolved distributions of core clock gene expression in livers of humans with obesity.** Livertranscriptomes were obtained by bulk RNA sequencing of liver biopsies from patients with obesity undergoing bariatric surgery and for which the exact daytime of sample collection was known. Biopsies were classified into three liver disease groups: “HN” (histologically normal), “NAFL” {benign steatosis only) or “NASH” (steatosis + inflammation) following a decision tree (see Fig.l). Each row of panels corresponds to a core clock gene. Left panels are violin plots with integrated boxplots comparing gene expression in morning (AM) and afternoon (PM) biopsies for each of the three liver disease groups. Statistics are Kruskal-Wallis tests followed by unpaired Wilcoxon post-hoc tests for AM-PM comparisons within each group (*p<0.05, **p<0.01, ***p<0.001). Central panels are dot plots with a lineartrendline of gene expression over daytime. Right panels are smoothed density plots of AM and PM gene expression distributions.

**Supplementary Figure S2.**
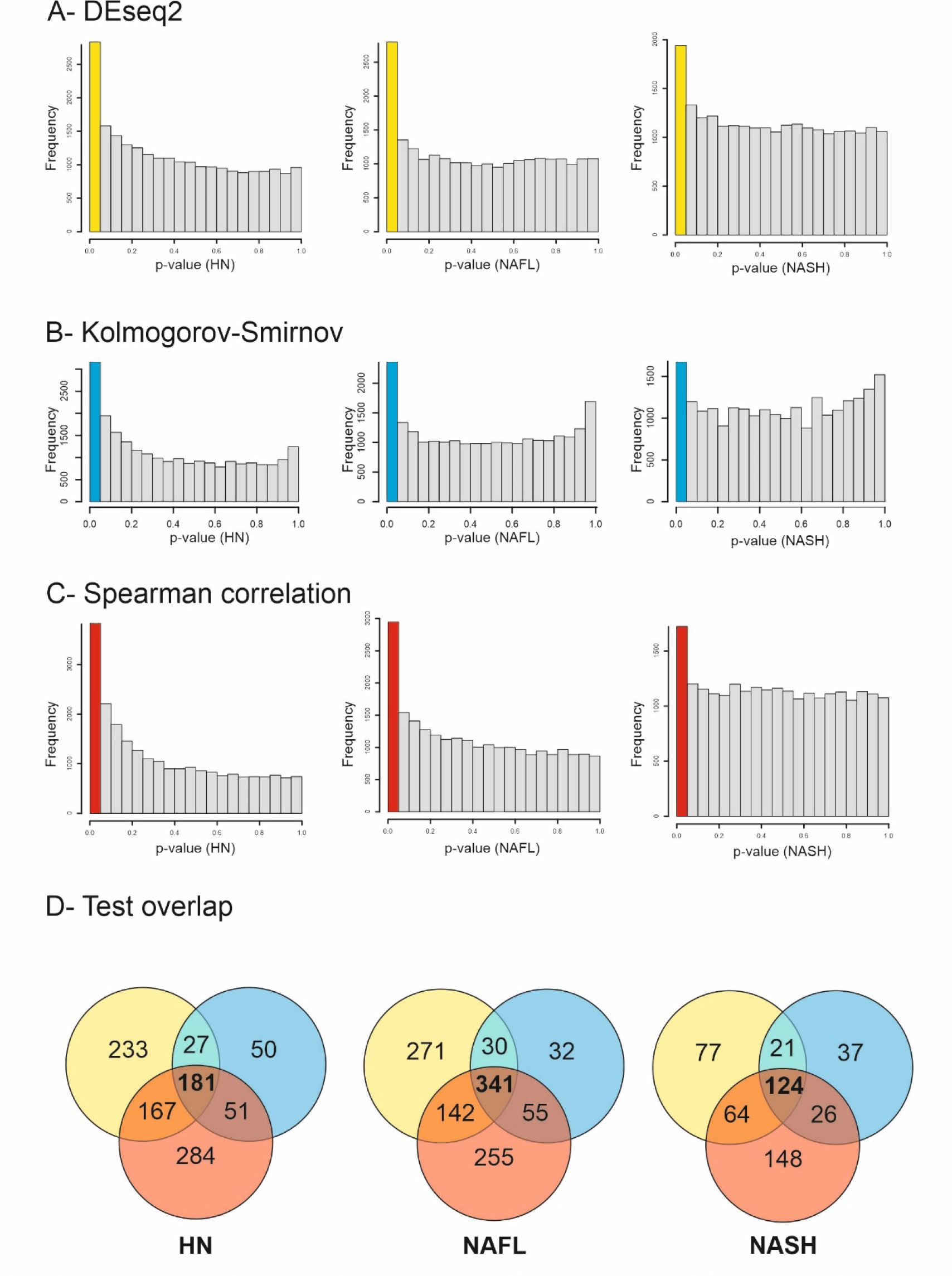
Comparison of the three independent statistical tests used to reveal daytime-dependency of gene expression. In absence of a full 24-hour time window necessary for standard, sinusoidal function-based methods to analyze circadian rhythmicity, we turned towards three different but complementary approaches to identify time-dependent genes (TDGs) in liver: A: DEseq2 based on means comparison of morning (AM) versus afternoon (PM) gene expression; B: Kolmogorov-Smirnov (KS) test to compare shapes of AM and PM gene expression distributions. C: Partial Spearman correlation between gene expression and daytime, A-C: Each row of panels corresponds to one test, with the three first panels displaying frequency histograms of uncorrected p-values for each liver disease group (“HN”, histologically normal; “NAFL", benign steatosis only; “NASH", steatosis + inflammation). D: Distribution of significant (FDR<O.l) TDGs from the corresponding test among groups. Venn diagrams (display the overlap between significant (FDR<O.l) TDGs from the three independent tests and were represented separately for each group.

**Supplementary Figure S3.**
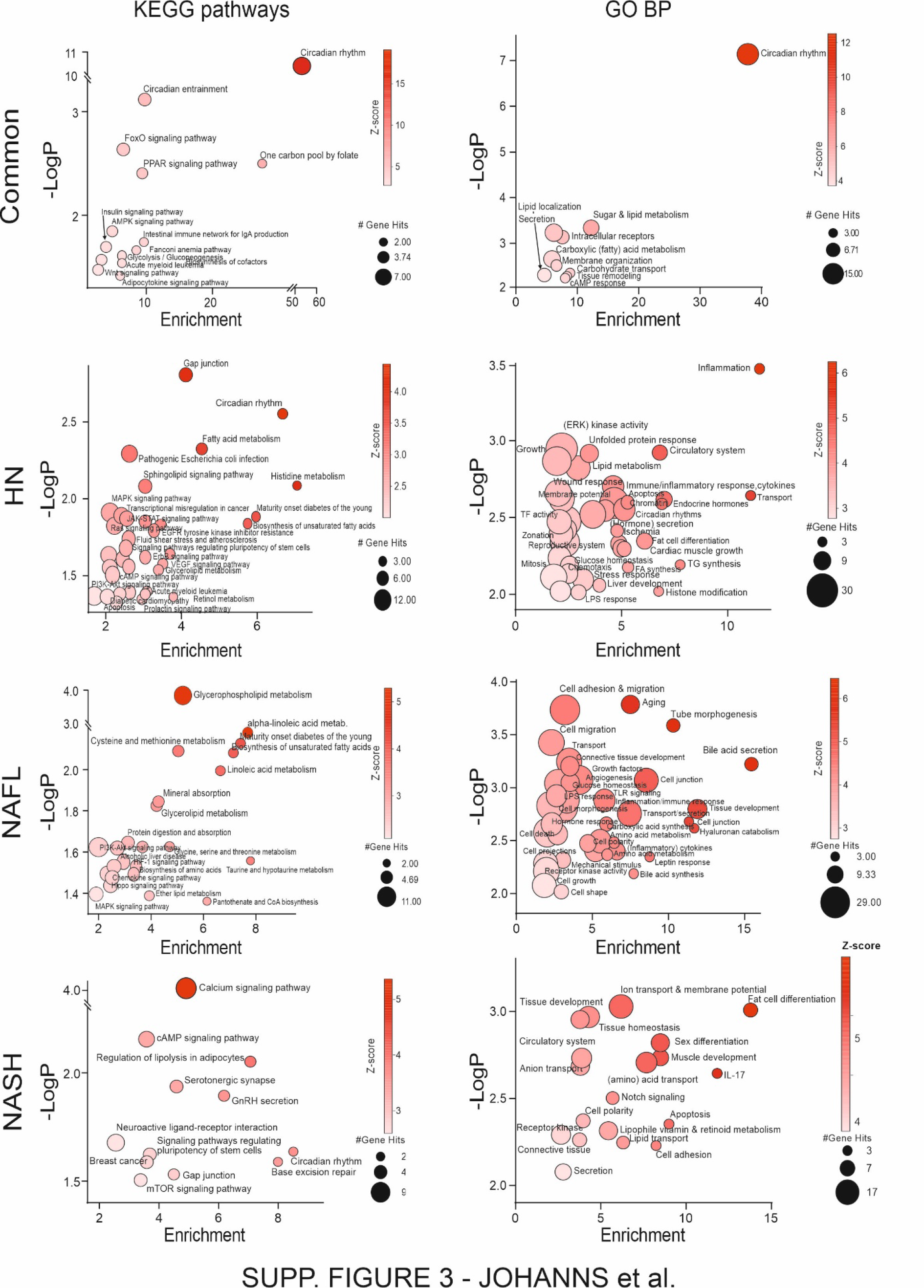
**(corresponding to main** **Figures 2C** **& 3). Biological term enrichment analysis of time-dependent genes (TDGs) for gene ontology biological process (GO-BP) and KEGG pathway terms.** Lists of significantly varying TDGs (Fisher FDR <0.01 and absolute AM-PM gene expression fold-change >1.2) were extracted as detailed in main Figure 2 for each liver disease group (“HN", histologically normal; “NAFL", benign steatosis only; “NASH", steatosis + inflammation). The online gene enrichment analysis platform Metascape was used to calculate enrichment in KEGG pathways (left panels) and GO-BP terms (right panels) for common or group- specificTDGs. Clusters were generated by Metascape and were manually renamed for visualization purpose and the lowest p-value from each cluster is displayed. Results are displayed as bubble plots, where the color gradient corresponds to the p-value of the enrichment and the size to the number of gene hits for each cluster.

**Supplementary Figure S4.**
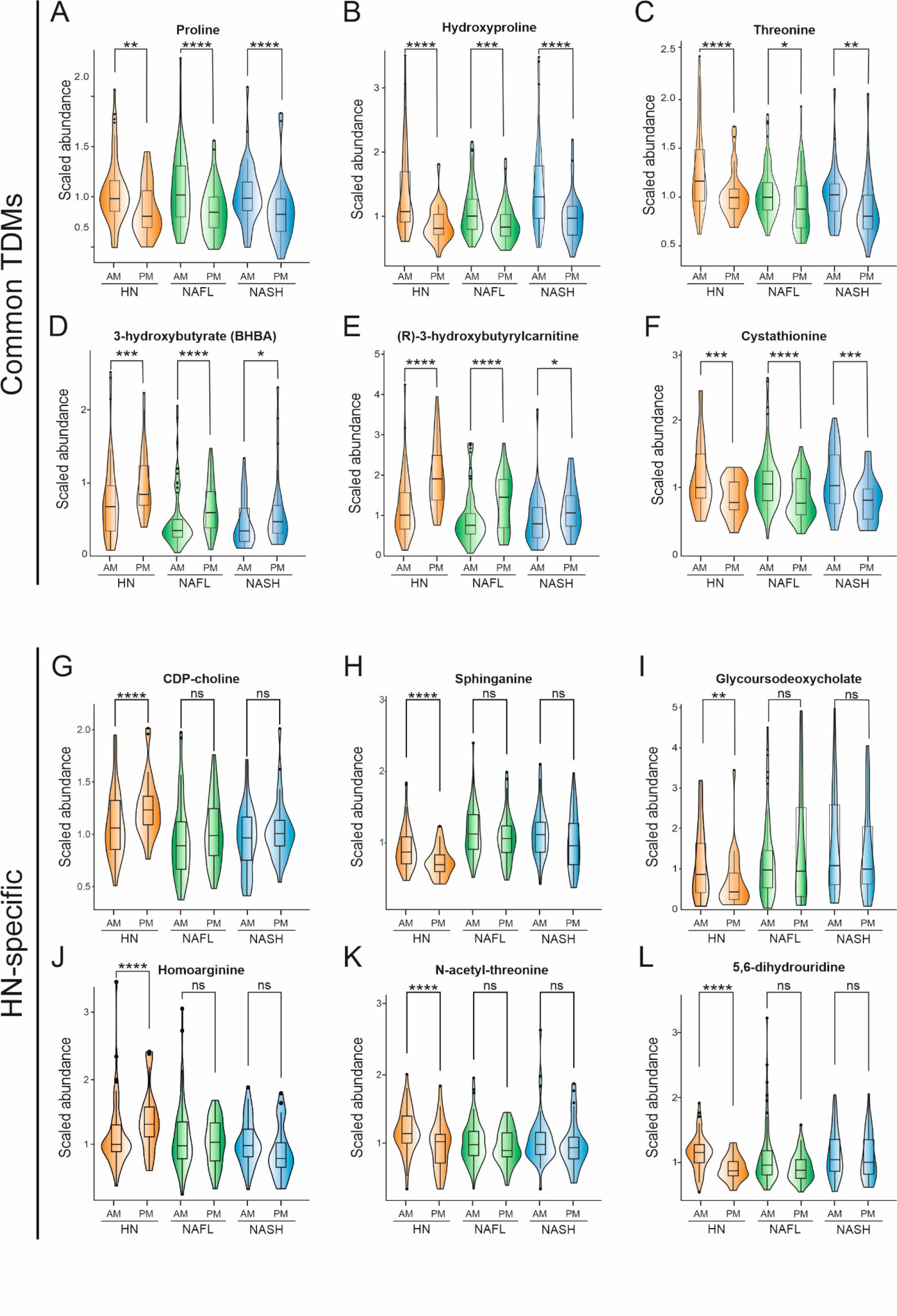
**Example of distributions of time-dependent liver tissue metabolites (TDMs) altered or not in NAFLD.** Liver tissue metabolites from human liver biopsies with known timing were obtained by multi-mode LC-MS. Biopsies were classified into three liver disease groups: “HN” (histologically normal), “NAFL” (benign steatosis only) or “NASH” (steatosis + inflammation) following a decision tree (see main Fig.l). TDMs were identified using the method detailed in main Figures 5 & 6 and were found to be either common to all 3 groups or specific to the HN group. Graphs are violin plots with integrated boxplots comparing metabolite abundance in morning (AM) and afternoon (PM) biopsies for each of the three liver disease groups. Statistics are Kruskal-Wallis tests followed by unpaired Wilcoxon post-hoc tests for AM-PM comparisons within each group (*p<0.05, **p<0.01, ***p<0.005, ****p<0.001).

**Supplementary Figure S5.**
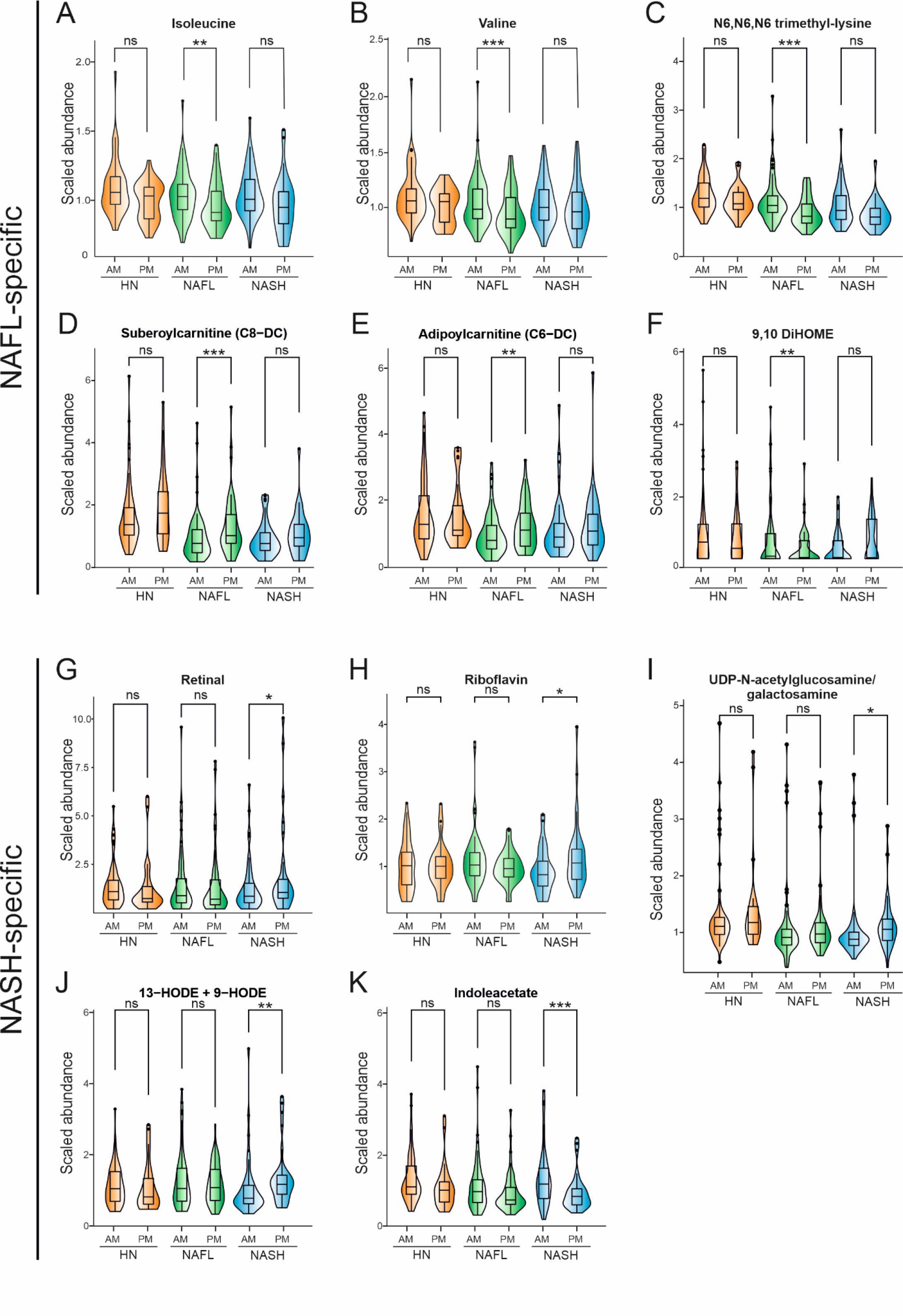
**Example of distributions of time-dependent liver tissue metabolites (TDMs) altered or not in NAFLD.** Liver tissue metabolites from human liver biopsies with known timing were obtained by multi-mode LC-MS. Biopsies were classified into three liver disease groups: “HN” (histologically normal), “NAFL” (benign steatosis only) or “NASH” (steatosis + inflammation) following a decision tree (see main Figure 1). TDMs were identified using the method detailed in main Figures 5 & 6 and were found to be specific to either the NAFL or NASH groups as indicated. Graphs are violin plots with integrated boxplots comparing metabolite abundance in morning (AM) and afternoon (PM) biopsies for each of the three liver disease groups. Statistics are Kruskal-Wallis tests followed by unpaired Wilcoxon post-hoc tests for AM-PM comparisons within each group (*p<0.05, **p<0.01, ***p<0.005, ****p<0.001).

